# Structure of an oxygen-induced tubular nanocompartment in *Pyrococcus furiosus*

**DOI:** 10.1101/2025.03.19.643997

**Authors:** Wenfei Song, Jan Fiala, Ioannis Skalidis, Pascal Albanese, Constantinos Patinios, Marten L. Chaillet, Servé W.M. Kengen, Richard A. Scheltema, Stuart C. Howes, Albert J.R. Heck, Friedrich Förster

**Author notes:** Equally contributed (Jan Fiala, Ioannis Skalidis). Univ. Grenoble Alpes, CNRS, INRAE, CEA, LPCV, INSERM, UMR BioSanté U1292, 38000 Grenoble, France.

## Abstract

Reactive oxygen species (ROS) pose a significant threat to biological molecules, prompting organisms to develop systems that buffer oxidative stress and contain iron, which otherwise amplifies ROS production. Understanding oxidative stress responses requires identifying the key proteins involved and their cellular organization. Here, we combined proteomics and cryo-EM to investigate the response of the anaerobic hyperthermophilic archaeon *Pyrococcus furiosus* to oxygen exposure. Proteome analysis revealed a significant upregulation of the oxidoreductase Rubrerythrin (Rbr) under oxidative stress. Cryo-electron tomograms showed the formation of prominent oxidative stress-induced tubules (OSITs). Single-particle cryo-EM and mass spectrometry of enriched OSITs identified them as stacked rings of Rbr homotetramers. The 3.3 Å structure demonstrates that rubredoxin-like domains mediate homotetramer assembly, suggesting that their oxidation drives OSIT formation. Within OSITs, we discovered virus-like particles formed by a ferritin-like/encapsulin fusion protein with iron hydroxide cores, uncovering a sophisticated organelle that protects *P. furiosus* from ROS through advanced compartmentalization.

## Introduction

Exposure of cells to oxygen causes oxidative stress due to formation of reactive oxygen species (ROS) such as superoxide and hydrogen peroxide, which can damage biological molecules^1^. Inside cells, ferric iron catalyzes the generation of further ROS from hydrogen peroxide through Fenton’s reaction, which amplifies oxidative stress and the need to respond to it. While aerobic organisms constitutively maintain antioxidant enzymes, anaerobic organisms feature redox-sensitive response systems, typically tailored to their environmental conditions. In shallow marine vents, the anaerobic *Pyrococcus furiosus* encounters oxygen when sulfide-rich vent fluid meets oxygenated seawater and possesses mechanisms to withstand transient oxygen exposure.

The primary response of anaerobes to superoxide is centered around the oxidoreductase superoxide reductase (SOR), which can catalyze superoxide reduction using rubredoxin as a donor^2^. The *P. furiosus* genome encodes several other oxidoreductases that may play additional, less understood roles in the oxidative stress response. In addition to these enzymatic systems, some bacteria and archaea evolved proteinaceous compartments that may sequester harmful metabolic products^3^. Such virus-like particles (VLPs), which measure 24 - 42 nm in diameter, are prevalent in many prokaryotes. The icosahedral VLPs first described in *P. furiosus* are composed of so-called encapsulin proteins^4,5^. In their core, the VLPs store ferrihydrite, which is thought to be an important component of the oxidative stress response^6^. The Fe^3+^ ions in the ferrihydrite precipitates are not reactive, which protects cells from ROS generation by Fenton’s reaction.

Current knowledge of the different molecular components of the oxidative stress response machinery of anaerobic organisms, and their interactions, remains incomplete. In this study, we combined mass spectrometry and different cryo-electron microscopy (EM) techniques to investigate the changes of the *P. furiosus* proteome and to determine its 3D intracellular organization under oxidative stress. Remarkably, we found that rubrerythrin (Rbr) tetramers assemble into oxidative stress-induced tubules (OSITs) upon oxygen exposure. These OSITs encapsulate VLPs, forming OSIT-VLPs complexes that link oxidoreductases and iron compartmentalization in mitigating redox stress.

## Results

### Rubrerythrin levels increase upon oxidative stress

To explore the proteome response to oxygen *P. furiosus* cells initially grown anaerobically on maltose as an energy and carbon source for different time periods (11, 15, 18, and 33 h) were exposed to an oxygen-containing atmosphere for approximately 30 min. This culture time includes the transition from the exponential growth phase into the stationary phase at approximately 18 h (Supplementary Fig. 1a). We collected a fraction from the cultured cells and utilized a label-free quantification (LFQ) mass spectrometry (MS) based proteomics approach to assess changes in protein abundance levels upon oxygen exposure.

Among all detected proteins, Rubrerythrin (Rbr) (corresponding to gene PF1283) consistently exhibits the highest increase in abundance across all time points when cells were exposed to an oxygen-containing atmosphere. Under aerobic conditions, Rbr levels are 3 to 13 times higher compared to anaerobic conditions (Fig. 1a-d). Notably, Rbr is the only protein that shows upregulation at every time point examined. In contrast, encapsulin (Enc, PF1192), another protein involved in the oxidative stress response, did does show significant variation in abundance under any of the conditions explored (Fig. 1f).

**Fig. 1.**
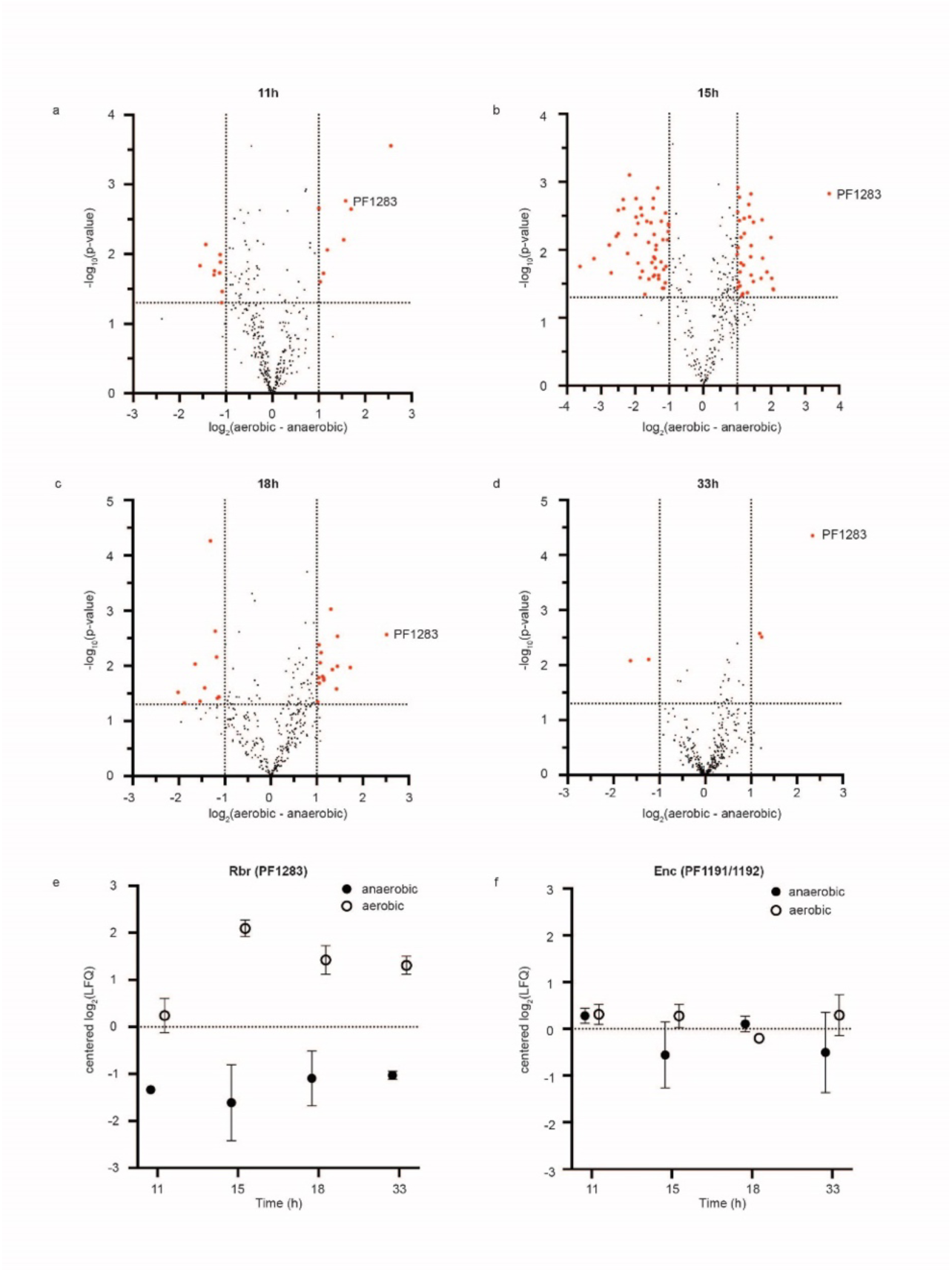
*P. furiosus* proteome response to oxidative stress. Volcano plots depicting the Rbr (PF1283) expression level change (based on Label-Free Quantification, LFQ) under aerobic *versus* anaerobic conditions. In panels (a-d), red dots denote statistically significantly up- and down-regulated proteins at 11h, 15h, 18h, and 33h of culture growth time. The horizontal dotted line indicates significance threshold of 0.05 for FDR-corrected p-values determined by Student’s t-test and a permutation test. Vertical lines represent fold change (FC) threshold set to ≥1. (e, f) Change in the expression profiles of rubrerythrin and encapsulin under anaerobic and aerobic conditions. Volcano plots data is listed in Supplemental Table 2.

Rbr is an oxidoreductase found in anaerobic bacteria and archaea. It has been shown to function as a hydrogen peroxide reductase across different species, while other redox functions such as ferroxidase and superoxide reductase (SOR) have also been reported^7^. Rbr has a ferritin-like (FL) di-iron domain and a mono-Fe rubredoxin-like (RubL) domain^4,8^ . In the cells, our mass spectrometric analysis also identified eight additional FL-domain-containing proteins that do not possess a RubL domain. However, the levels of these FL-domain-containing proteins do not vary substantially under anaerobic *versus* aerobic conditions (Supplementary Fig. 1b). This suggests that Rbr plays a distinct and critical role in the oxidative stress response, likely due to the presence of its unique RubL domain, setting it apart from its FL-domain-containing counterparts.

### Rbr forms oxidative-stress-induced tubules

To analyze how oxidative stress changes the intracellular three-dimensional (3D) organization of macromolecules, we performed cryo-electron tomography (ET) of whole *P. furiosus* cells. As a baseline, we first imaged the cells in their preferred anaerobic environment. In the tomograms, ribosomes are readily discernable in the cytosol, as expected for intact cells (Supplementary Fig. 2a and Supplementary Video1). Consistent with the abundance of Enc in the cells, many tomograms also display virus-like particles (VLPs), which have highly dense lumens.

We then imaged *P. furiosus* cells that were exposed to air after anaerobic growth. To our surprise, many tomograms of these stressed *P. furiosus* cells reveal tubular nanocompartments, which have not been observed in any of the tomograms of anaerobically grown cells (Fig. 2a, 2b, Supplementary Fig. 2b, 2c, Supplementary Video2-5). Due to its emergence upon oxidative stress and its tubular shape we refer to this nanocompartment as an oxidative-stress-induced tubule (OSIT). In the OSIT-containing tomograms, OSIT particles measure approximately 45 nm in diameter and vary strongly in length (40-490 nm, in tomograms covering approximately 900 nm x 900 nm) (Supplementary Fig. 2d).

**Fig. 2.**
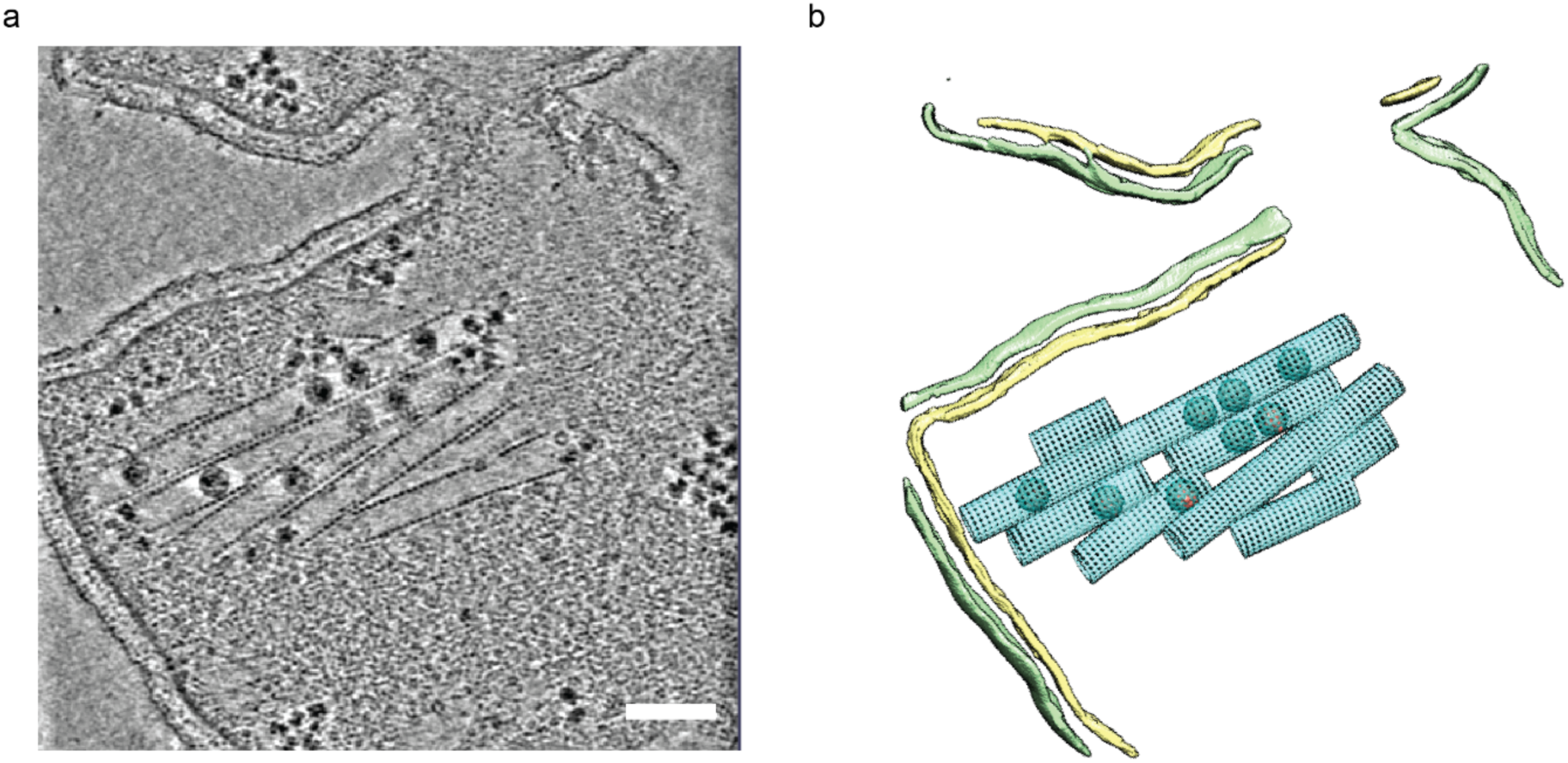
Oxidative stress-induced tubules (OSITs) formation upon oxidative stress. (a) Cryo-tomogram slice of *P. furiosus* cell grown under anaerobic conditions after 33 h cultivation then exposed to oxygen containing environment. Scale bar: 100 nm. (b) 3D isosurface rendering of segmented cell shown in a. cyan: OSIT, pink: VLPs, yellow: cell membrane, green: S-layer.

To identify the molecular composition of these OSITs, we enriched OSIT from air-exposed, lysed *P. furiosus* cells using sucrose gradient centrifugation. To identify OSIT-enriched fractions they were screened using negative stain EM. Those fractions showing abundant OSITs were further analyzed using proteomics, which identified Rbr as the most abundant protein (Supplementary Fig. 3a). To confirm the identity of the primary OSIT component and to obtain high-resolution structural details, we subjected the OSITs to cryo-EM analysis. Cryo-ET combined with sub-tomogram analysis of OSIT segments revealed that the OSITs have diameters ranging from approximately 46 nm to 55 nm and consist of rings stacked at intervals of approximately 8 nm. Classification analysis indicated that these rings are composed of 16 to 19 subunits, with the most common configuration being 17 subunits, resulting in an OSIT diameter of approximately 48 nm (Supplementary Fig. 3b).

To determine a more precise OSIT spatial architecture, we acquired cryo-EM projection images, which we firstly analyzed by helical reconstruction. Focusing on the most abundant OSIT class with 17 subunits per ring the overall density of the OSIT was first determined to a resolution of approximately 11 Å by aligning and averaging OSIT segments of approximately 10 stacked rings (98.4 nm) in a reconstruction volume (Supplementary Fig. 4, 5). To further increase the resolution, which may be limited by variations of ring assembly and stacking, we focused the alignment on a 3×3 subunit patch, resulting in a 7.4 Å resolution map. Consistent with this resolution assessment, the resulting map revealed α-helical secondary structure features. The reconstruction shows that each ring subunit is a homotetramer. Finally, we used unconstrained SPA of 2×2 patches to obtain an EM density map with a resolution of approximately 3.3 Å (Fig. 3a, Supplementary Fig. 6a, Supplementary Table 1). The map allows unambiguous positioning of Rbr X-ray crystallographic models^9^ and further refinement of the atomic model (Supplementary Fig. 6b).

**Fig. 3.**
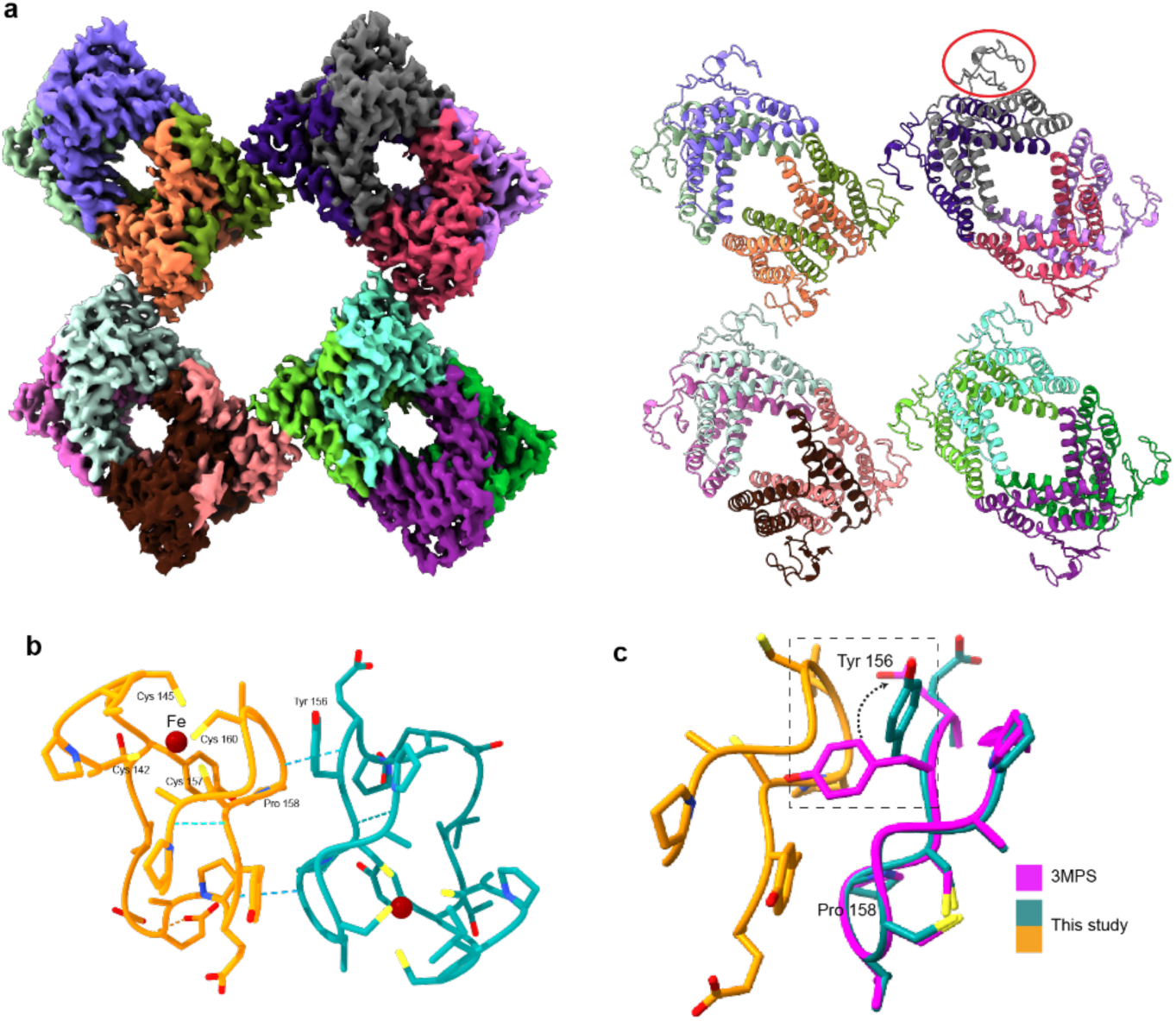
RubL domain contributes to OSIT formation. (a) Left: cryo-EM density map of 2×2 Rbr tetramer patch in OSIT, colored by fitted subunit. Right: atomic model of four Rbr tetramers built into cryo-EM map. A RubL domain is marked by a circle. (b) Interface of neighboring tetramers formed by RubL domains of Rbr monomers from adjacent tetramers. Dotted lines indicate hydrogen bonds.(c) Comparison of our atomic model in OSIT with crystal structure of oxidized Rbr (3MPS). Arrow indicates repositioning of Tyr156 side chain.

### RubL domain drives OSIT formation

The cryo-EM map reveals that the inter-tetramer contacts within each ring and the longitudinal contacts between the rings are both formed by adjacent RubL domains (Fig. 3a). The core of each RubL domain is a cluster of four cysteines (Cys 142, 145, 157, 160), which coordinate an Fe ion [Fe(SCys)_4_]. This cluster appears intact in the cryo-EM map and it seems to stabilize the Rbr interface (Fig. 3b): two of the involved cysteine side chains (Cys 157, 160) belong to the loop (residues 155-163) that constitutes the tetramer-tetramer interface. This loop is stabilized by the Fe coordination and interacts with its counterpart from the adjacent Rbr tetramer via two twin hydrogen bonds that form between Tyr 156 and Pro 158 (Fig. 3b).

Comparison of the OSIT atomic model to X-ray crystallographic structures of different oxidation states of the Rbr FL domain allows for assessing the oxidation state (Fig. 3c). Among the different states, the peroxide-bound oxidized Rbr crystal structure (PDB ID: 3MPS) is essentially identical to the OSIT model. Thus, the FL-domain adopts the oxidized conformation in the context of Rbr. The different positioning of the Tyr-156 side-chain in the crystal structure may be due to the fact that Rbr was in its reduced form when crystals were formed^9^, which might have constrained conformational changes upon oxidation and prevented OSIT formation.

The close agreement between the crystal structure of fully oxidized Rbr and the OSIT map suggests that oxidation is responsible for OSIT formation. To support this notion biochemically, the enriched OSIT fraction was treated with 5mM DTT, a reducing agent. Negative stain EM shows that this treatment results in the dissociation of the OSIT nanocompartments (Supplementary Fig. 7), indicating that reduction reverses OSIT formation.

### OSIT encapsulates virus like particles under oxidative stress

After Rbr, the second most abundant iron-binding protein in the *P. furiosus* proteome is Enc. Encapsulins form icosahedral virus-like particles (VLPs) by assembling 180 Enc copies, with a structure similar to the fold of the Escherichia virus HK97 protein^4^. VLPs are common in many prokaryotes and often encapsulate FL-proteins as cargo, which enables VLPs to bind and sequester ferric iron^8^. This sequestration of iron is thought to protect organisms from oxidative damage^6^. In *P. furiosus*, the genes encoding Enc (PF1192) and the FL domain (PF1191) were previously suggested to give rise to a single FL-Enc fusion protein^5^, which we could confirm by proteomics (Supplementary Fig. 8, 9, 10). In the cellular tomograms, the VLPs have a pronounced high density in their lumens, consistent with the sequestration of iron hydroxide.

To our surprise, the tomograms of stressed *P. furiosus* cells reveal VLPs encapsulated inside the OSITs (Fig. 2a, 2b, Supplementary Fig. 2b, 2c, Supplementary Videos 2-5). The VLPs distribute somewhat irregularly in the OSITs, with some tubules completely devoid of VLPs. These observations suggest that VLPs and OSITs assemble independently, with VLPs either being encapsulated during the OSIT assembly or diffusing into the OSITs after both structures have fully formed.

### Rbr knockout abolishes OSIT formation and shows magnesium and calcium granules

To validate that Rbr is required for OSIT formation, we generated a *P. furiosus* Rbr knock-out (*Δrbr*) strain. Indeed, proteomics analysis of the *Δrbr* cells confirmed the absence of Rbr (Supplementary Fig. 9). Compared to the wildtype (WT) strain, the *Δrbr* strain has a slightly longer lag phase during growth under anaerobic conditions, while the growth rate at late stages increases beyond a level seen in the wildtype cells (Supplementary Fig. 1a).

Like in the wildtype cells under anaerobic conditions, none of the cryo-ET volumes of the *Δrbr* cells display OSITs, while we do observe VLPs. Different from the wildtype cells though, we did not detect any OSITs upon oxygen exposure in any of the tomograms from the *Δrbr* cells (Supplementary Video 6).

An unexpected characteristic of the *Δrbr* strain under aerobic conditions is the formation of dense granules inside the cell, which are clearly visible in the cryo tomograms (Supplementary Fig.10a, Supplementary Video 6). To determine the elemental composition of these granules, we used energy dispersive spectroscopy (EDX) analysis on the frozen-hydrated sample. The X-ray spectra from the granules within these cells revealed that magnesium, calcium, and phosphorus are their main inorganic constituents (Supplementary Fig. 10b-10f). This observation suggests that the Rbr knockout affects the cell’s metabolism in ways beyond the scope of this study.

## Discussion

In this study we combined cryo-ET imaging with proteomics analysis to study the response of *P. furiosus* to oxidative stress. Archaea, including *P. furiosus*, often incorporate iron as a cofactor, likely due to the iron-rich environments they inhabit^10^. A major challenge for *P. furiosus* under aerobic conditions is the formation of ROS via Fenton’s reaction, which is catalyzed by Fe^2+^. Hence, it is vital for the cell to maintain low levels of free Fe^2+^ by oxidizing and sequestering it.

Rubrerythrin (Rbr) is the most uniquely and substantially upregulated protein in *P. furiosus* in response to oxidative stress. Our most surprising finding is that during oxygen exposure, Rbr assembles into OSITs, which can encapsulate VLPs. The high density in the lumens of the VLPs in our cellular tomograms indicates a high concentration of inorganic precipitates at these sites, which interact much stronger with electrons than organic material. This observation is in line with the strong accumulation of ferrihydrate, which has previously been studied in the bacterium *Myxococcus xanthus*^6^. Genetic studies in this organism confirmed that VLPs contribute to viability of cells exposed to oxygen, likely by protecting them from the toxic products of the Fenton reaction^6^.

### Oxidative stress induced formation of OSITs

Enzyme oligomerization, often in the form of large filaments, has recently emerged as a common mechanism to regulate their function, in particular in prokaryotes ^11^. *P. furiosus* Rbr is an oxidoreductase with two redox domains. Its di-iron FL domain has been shown to have peroxidase activity ^12^, while the function of the mono-iron RbrL domain is less clear. Already two decades ago, the striking tendency of Rbr to aggregate when exposed to oxygen was reported ^12^. We explain this phenomenon by the formation of OSITs upon oxidative stress, suggesting that the previously described aggregates were possibly Rbr tubules.

The quaternary structure of the OSITs suggests that the RbrL domain is key to OSIT formation because it forms all inter-tetramer contacts. The RbrL domain is only present in anaerobic archaea, while Rbr from aerobic archaea does not feature this domain^13^. This suggests that Rbr from aerobic archaea, which are fully adapted to oxygen, have lost the capability to form OSITs. We propose that RbrL may act as a redox potential dependent molecular switch, which controls OSIT formation.

Indeed, upon the introduction of a reducing agent (DTT) we observed a complete disassembly of OSITs. Mechanistically, this observation, along with the structural data presented in this work, are indications that the [Fe(SCys)_4_] cluster contained within the RubL domain of *P. furiosus* Rbr is indeed a molecular switch that triggers OSIT formation: upon oxidative stress, the Fe ion contained within the cluster will be oxidized, stabilizing the cysteines of the coordination site that are part of the polymerization interface loop (Cys 157, 160). This change is reversible, translating to a potential disruption of the interface and OSIT breakdown, upon loss of environmental oxidative stress. It is fascinating that subtle structural changes of Rbr induced by oxidation trigger a higher-order protein organization in response to environmental stress.

### Functional interaction of Rbr and Enc to combat oxidative stress

When *P. furiosus* cells encounter an oxygen-containing environment, oxygen can be oxidized to toxic superoxide produced, for example by electron carriers. Superoxide reductase (SOR) alleviates superoxide toxicity by reducing it to hydrogen peroxide, using rubredoxin as an intermediate and NADH as the ultimate electron donor^2^. We propose that the upregulation of Rbr and its assembly into OSITs play a key role in managing H_2_O_2_ in the cell (Fig. 4). Rbr’s FL di-iron domain functions as a peroxidase, reducing hydrogen peroxide to water by accepting electrons from the Fe(SCys)_4_ cluster in the RbrL domain^14,15^. Thus, Rbr may function as a scavenger for H_2_O_2,_ and upregulation increases capacity. Another consequence of the increased Rbr concentration is the binding of free Fe ions, which would otherwise promote the harmful Fenton reaction.

**Fig. 4.**
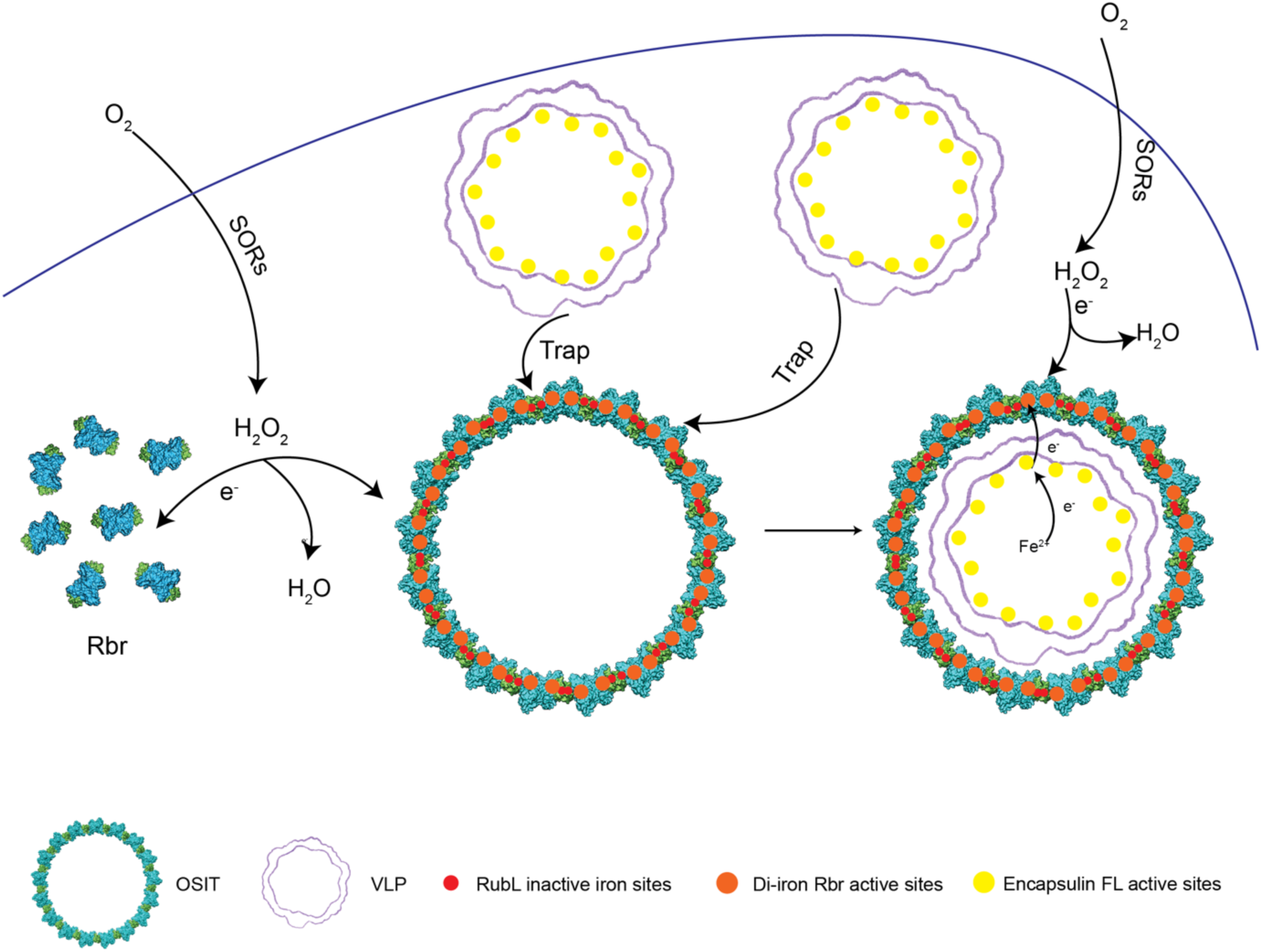
Proposed model for electron transfer upon oxidative stress. SOR firstly responds by reducing superoxide into hydrogen peroxide. The resulting hydrogen peroxide oxidizes Rbr, which triggers OSIT formation. OSITs trap VLPs and form VLP-OSIT super-complexes. VLPs can sequester Fe(OH)_3_ in their lumens to prevent catalysis of the harmful Fenton reaction. In the core of VLPs formed by FL-Enc, the FL domains, which assemble to dodecamers, can oxidize Fe^2+^ to Fe^3+^, which gets contained. The resulting electrons may flow across the VLP shell to Rbr’s di-iron FL domain, which has a higher redox potential. These electrons in turn can be accepted by further H_2_O_2_ molecules.

The oxidation of Rbr triggers the assembly into OSITs, which subsequently encapsulate FL-Enc VLPs. The FL domain of FL-Enc, which is localized within the VLPs similarly to FL proteins in VLPs from the bacterium *Myxococcus xanthus*^16^, catalyzes the oxidation of Fe^2+^ to Fe^3+^ and stores it as Fe(OH)_3_^17^. We speculate that Rbr, in its fully ferric di-iron state within OSITs, can act as an electron acceptor, enabling further H_2_O_2_ oxidation. Thus, the OSIT-VLP supercomplexes may function to oxidize intracellular peroxide during oxidative stress and to sequester iron.

## Conclusion

Using a combination of proteomics, cryo-ET, cryo-EM SPA, molecular modeling and functional studies we discovered assembly of Rbr into oxidative stress induced tubule (OSITs) in *P. furiosus*. Together with the VLPs encapsulated in their lumens, OSITs play a role in combatting oxidative stress. Like VLPs being widely conserved across prokaryotes, OSITs may also be common in anaerobes well beyond *P. furiosus* and may be exploited for biotechnological applications^19^.

## Acknowledgment

This work was supported by the Nederlandse Organisatie voor Wetenschappelijke Onderzoek (NWO) with research program TA 741.018.201 and the European Research Council under the European Union’s Horizon2020 Program (ERC Proof-of-Concept Grant Agreement 101113464–BENDER, to F.F.). We thank Marel Spoelstra for helping with preliminary experiments. We thank Hans Meeldijk for helping with EDX data acquisition and data analysis and the whole University Electron Microscopy Center for facility support. We thank Willem Noteborn for the support with data collection at NeCEN (National Roadmap for Large-Scale Research Infrastructure NEMI 184.034.014).

## Author contributions

W.S. and F.F. designed the project. W.S performed the cell culture, OSIT enrichment, cryo-EM and negative staining sample preparation, SPA, cryoET and EDX data acquisition and image analysis and DTT treatment of OSIT enrichment. I.S., W.S, and S.C.H. performed OSIT single particle analysis. I.S. built atomic models. P.A. and R.A.S. performed the proteomics analysis for enriched OSIT samples. C.P. and S.K. constructed the *Δrbr* strain. J.F. and A.J.R.H. performed the whole cell proteomics data collection and analysis for WT, *Δrbr* strain under anaerobic and aerobic conditions. M.C. helped with cryoET data processing. W.S and F.F. wrote the manuscript with input from all authors. R.S., A.J.R.H., and F.F. provided financial funding for the project.

## Declaration of generative AI and AI-assisted technologies

During the preparation of this work, the authors used ChatGPT 4.0 in order to improve clarity. After using this tool or service, the authors reviewed and edited the content as needed and take full responsibility for the content of the publication.

## Materials & Correspondence

The data that support this study are available from the corresponding author upon reasonable request. Raw tomography data will be deposited in EMPIAR and the single-particle reconstruction of OSIT is deposited in EMDB under accession number 53127 and the corresponding model in PDB under accession number 9QG1. Representative tilt series are deposited on EMPIAR under accession number 12630. MS data will be deposited in PRIDE, under accession number PXD050805.

**Supplementary Fig. 1.**
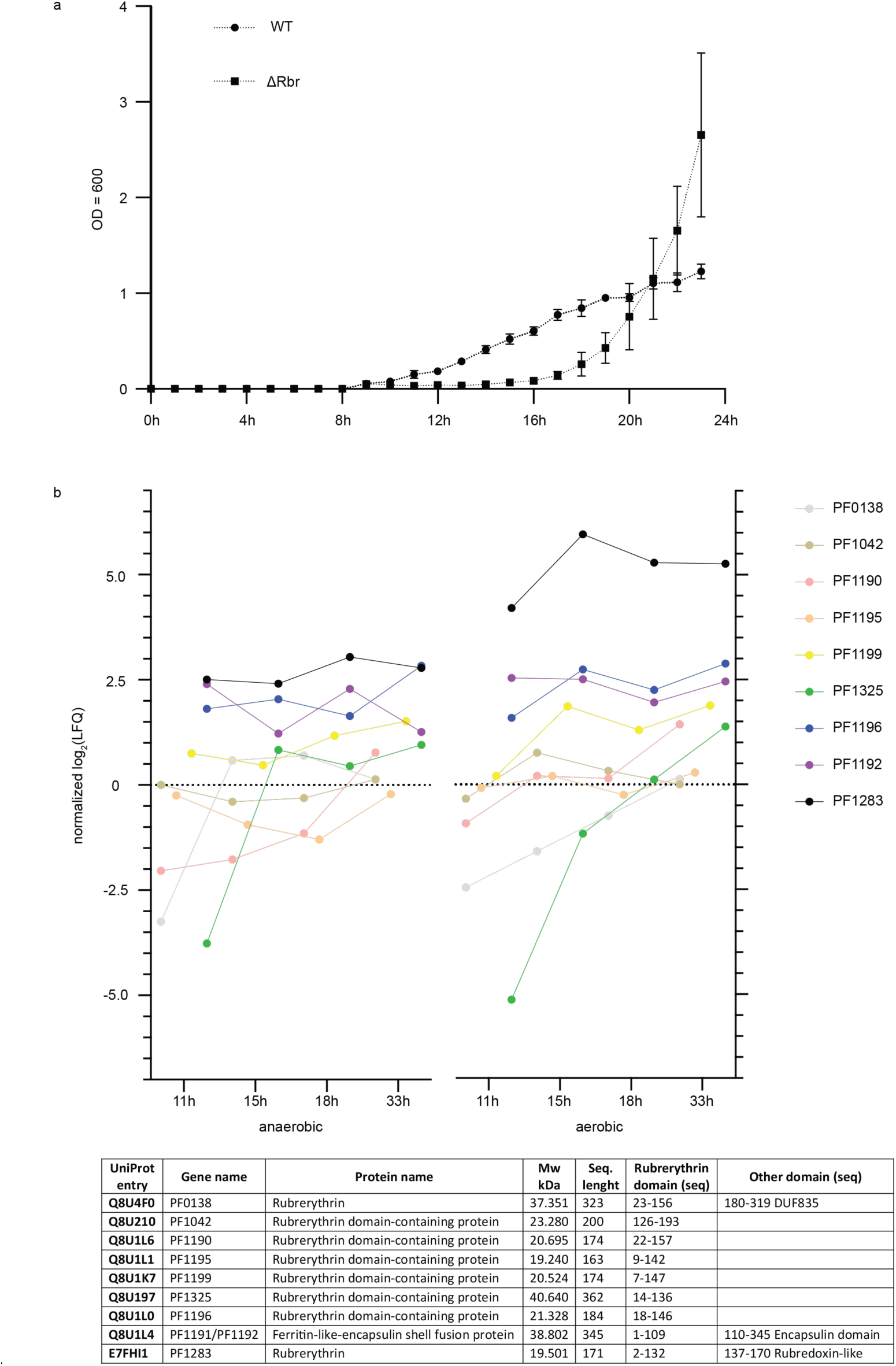
Response to oxidative stress in *P. furiosus* of Ferritin like domain containing proteins. (a) Growth curves of WT and *Δrbr P. furiosus* cells in anaerobic environment over 24 h. (b) 8 rubredoxin domain containing proteins detected and quantified by proteomics at 4 time points under anaerobic and aerobic conditions. Of all these proteins uniquely PF1283 is highly more abundant under aerobic conditions. The proteins are annotated according to their encoding genes.

**Supplementary Fig. 2.**
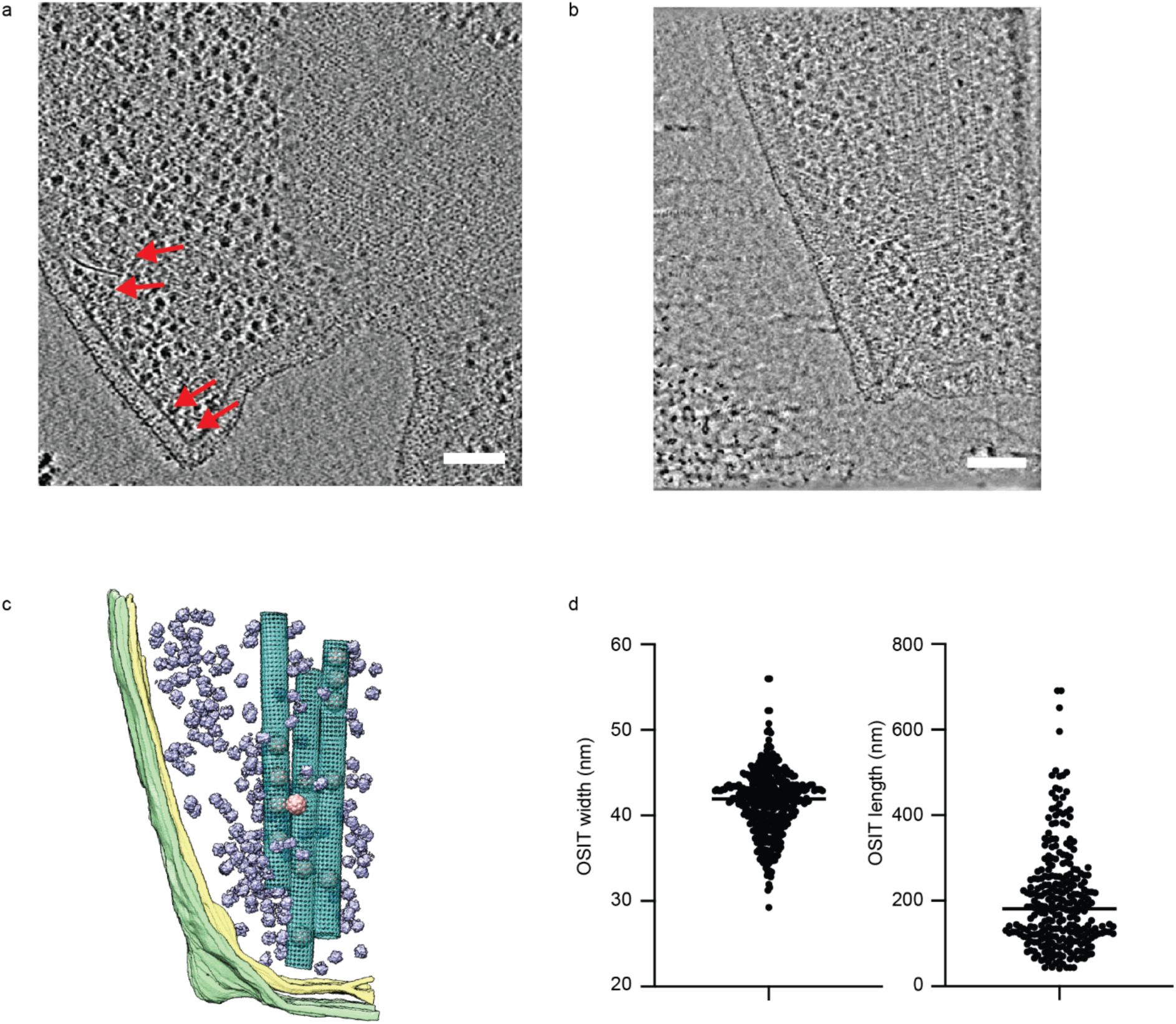
Cryo-tomogram slices of *P. furiosus* cells in response to oxidative stress. (a) Cryo-tomogram slice of *P. furiosus* cell under anaerobic conditions after 33 h cultivation. Along the z-direction the slices covers 1.73 nm. Red arrow: VLP. Scale bar: 100 nm. (b) Cryo-tomogram slice of *P. furiosus* cell under anaerobic conditions after 33 h cultivation then exposed to oxygen-containing environment. Scale bar: 100 nm. (c) 3D isosurface rendering of segmented cell shown in b. cyan: OSIT, pink: VLPs, purple: ribosomes, yellow: cell membrane, green: S-layer. Scale bar: 100 nm. (d) Analysis of OSIT width and length distribution in all tomograms with OSITs (N=421 OSIT, 120 tomograms).

**Supplementary Fig. 3.**
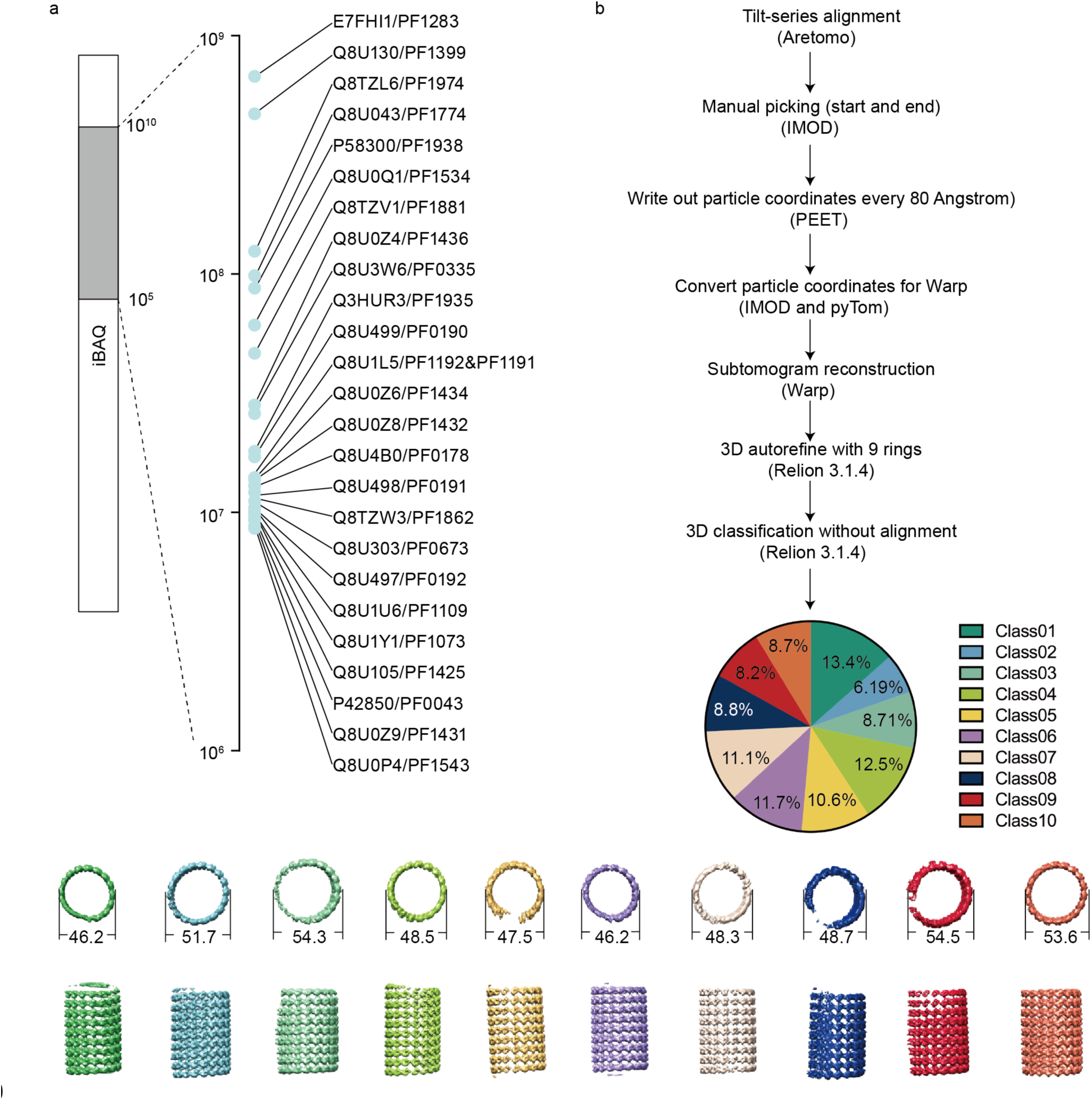
Identification of OSIT composition by MS and subtomogram averaging. (a) Intensity based quantification by label free proteomics shows the 25 most abundant proteins (Uniprot IDs) in the enriched OSIT sample, with Rbr being the most prominent (Supplementary Table 2). (b) OSIT subtomogram averaging indicates heterogeneity of the OSITs. The tomograms are aligned using Aretomo, particles were manually picked in IMOD, then converted the coordinates for Warp with a rise of every 80 angstrom. Subtomograms were reconstructed in Warp then applied 3D classification in Relion 3.1.4. The 3D classification shows 10 classes with various nanometers in diameter.

**Supplementary Fig. 4.**
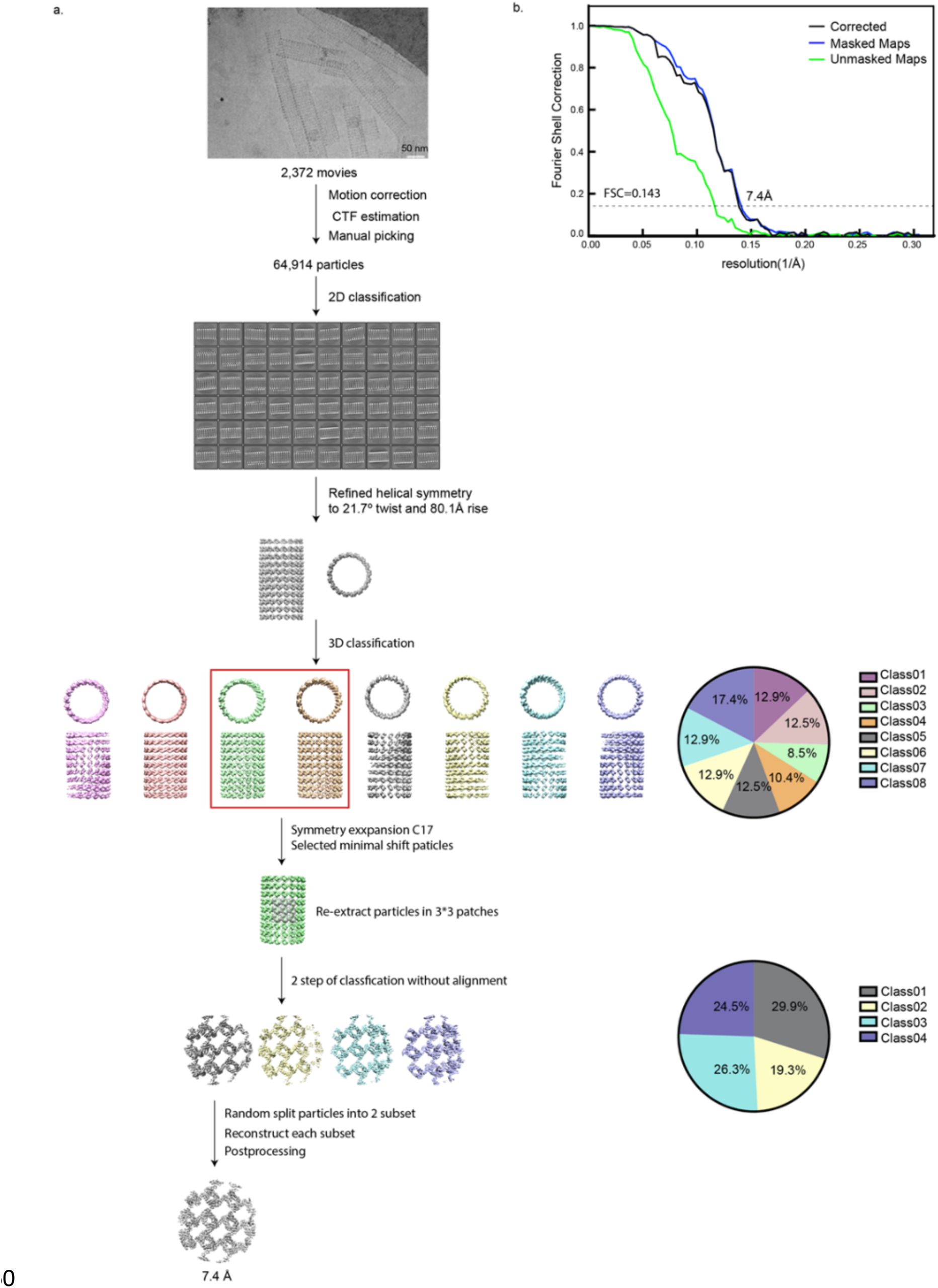
Helical reconstruction workflow for OSIT processing. (a) Workflow of the helical reconstruction detailed in Materials and Methods. Scale bar: 50nm. (b) Fourier shell correlation (FSC) curves of the 3D reconstruction of 3×3 patch density map.

**Supplementary Fig. 5.**
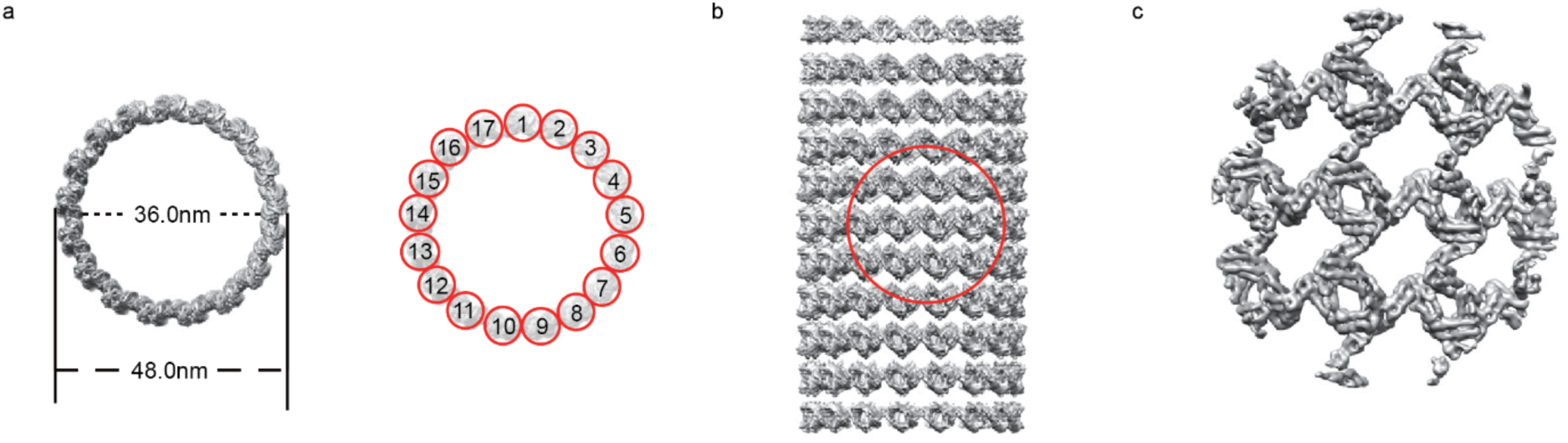
3D density map of Helical reconstruction of OSITs. (a) CryoEM density map of OSIT in top view. Left: CryoEM density map of OSIT in top view. Right: cross-section view of OSIT with subunits indicated as found in best-resolved class. (b) CryoEM density map of OSIT in side view. (c) cryoEM density of OSIT with enlarged view of circled area in (b).

**Supplementary Fig. 6.**
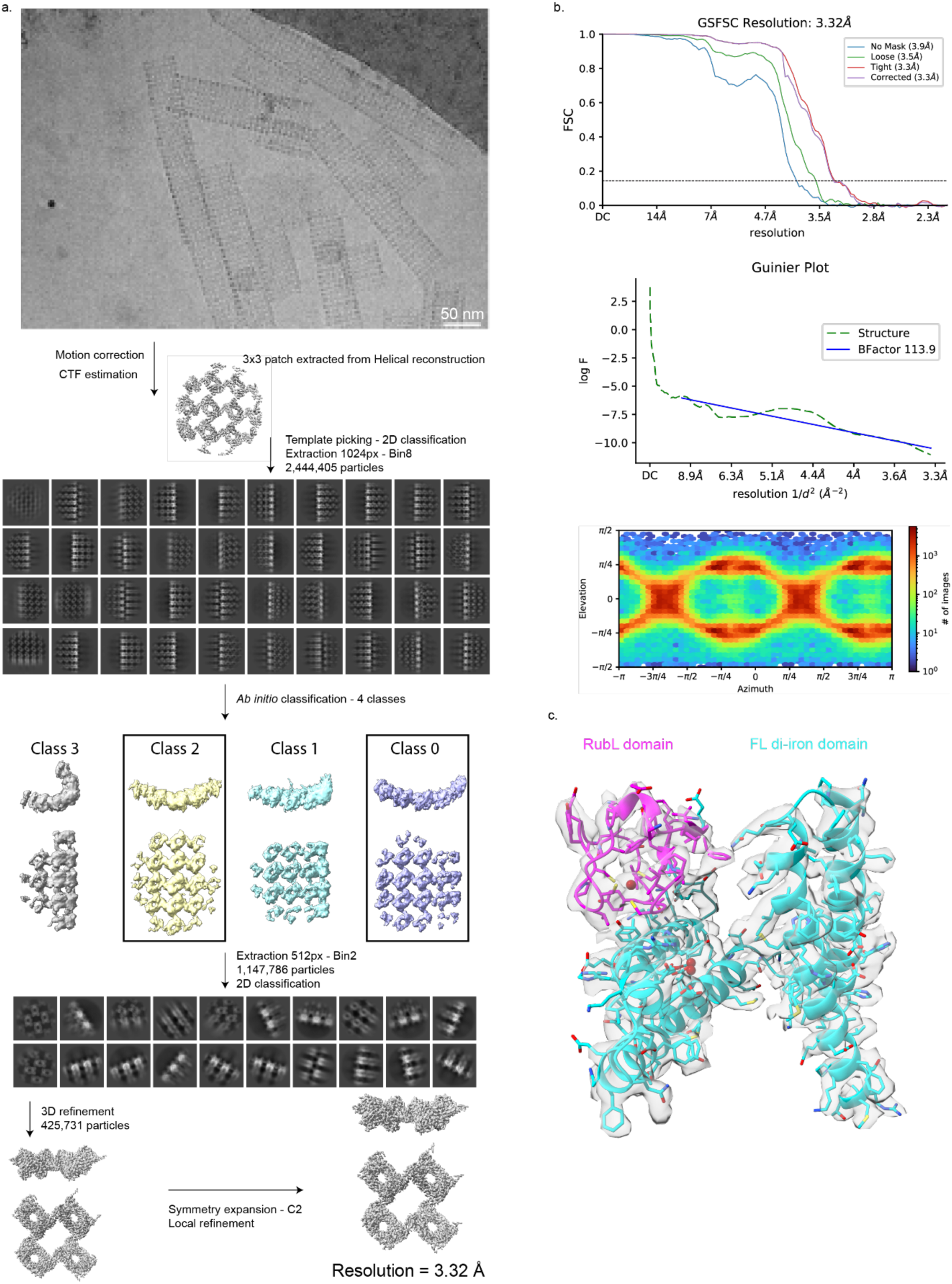
Cryo-EM single-particle analysis and validation. **(a)** Single-particle analysis workflow to reconstruct 2×2 patch Rbr density. **(b)** Fourier Shell Correlation (FSC), Guinier plot with fitted B-factor and particle orientation distribution plots for the final Rbr reconstruction. **(c)** Domain annotation and model/map fit of Rbr.

**Supplementary Fig. 7.**
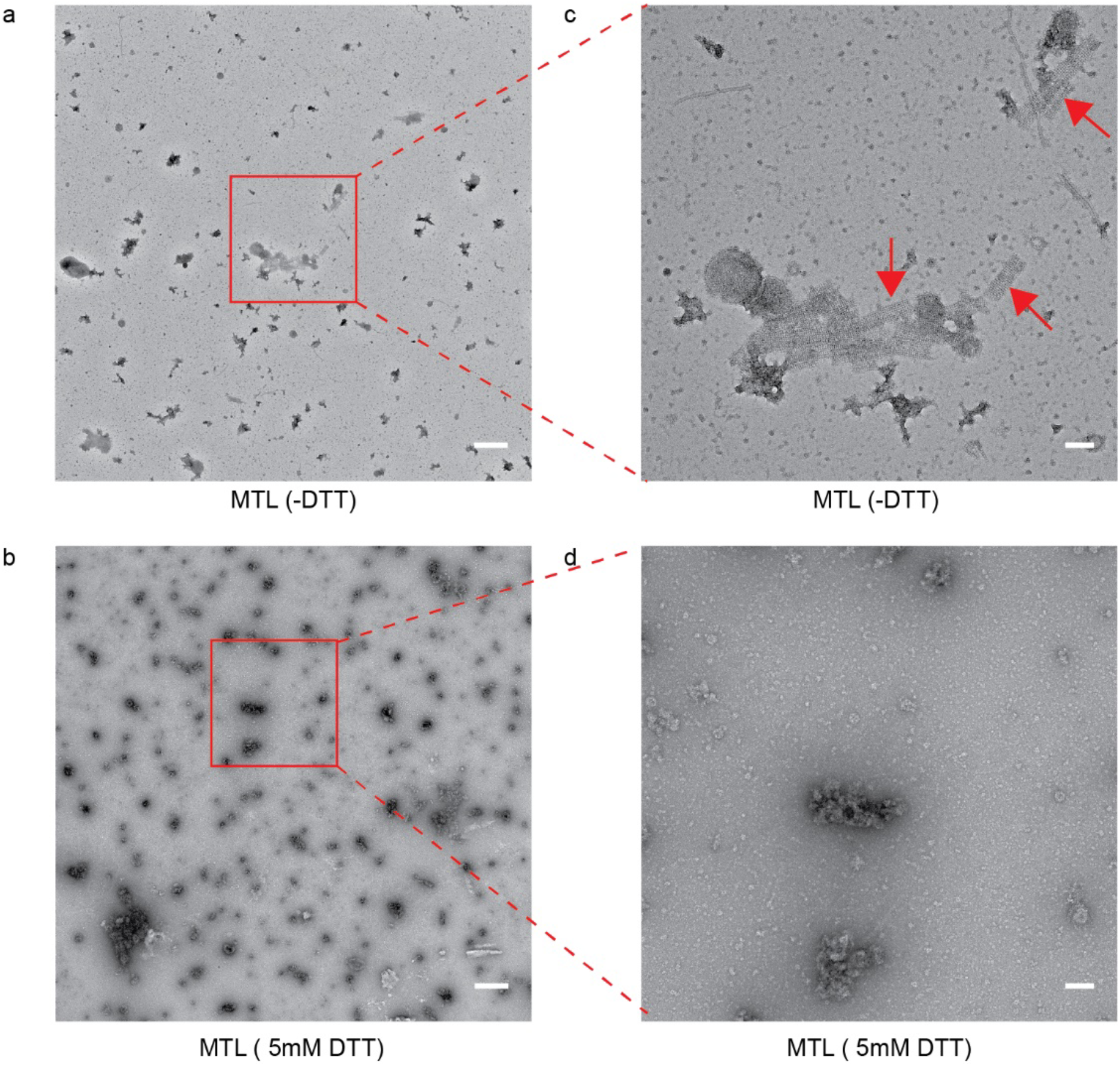
OSIT dissociation upon 5mM DTT treatment imaged by negative stain EM. (a) untreated enriched OSIT fraction and (b) enriched OSIT fraction treated with 5mM DTT at low magnification (scale bar: 500 nm). (c) enriched, untreated OSIT. The arrow indicates OSITs. (d) enriched OSIT treated with 5mM DTT at higher magnification (scale bar: 100nm).

**Supplementary Fig. 8.**
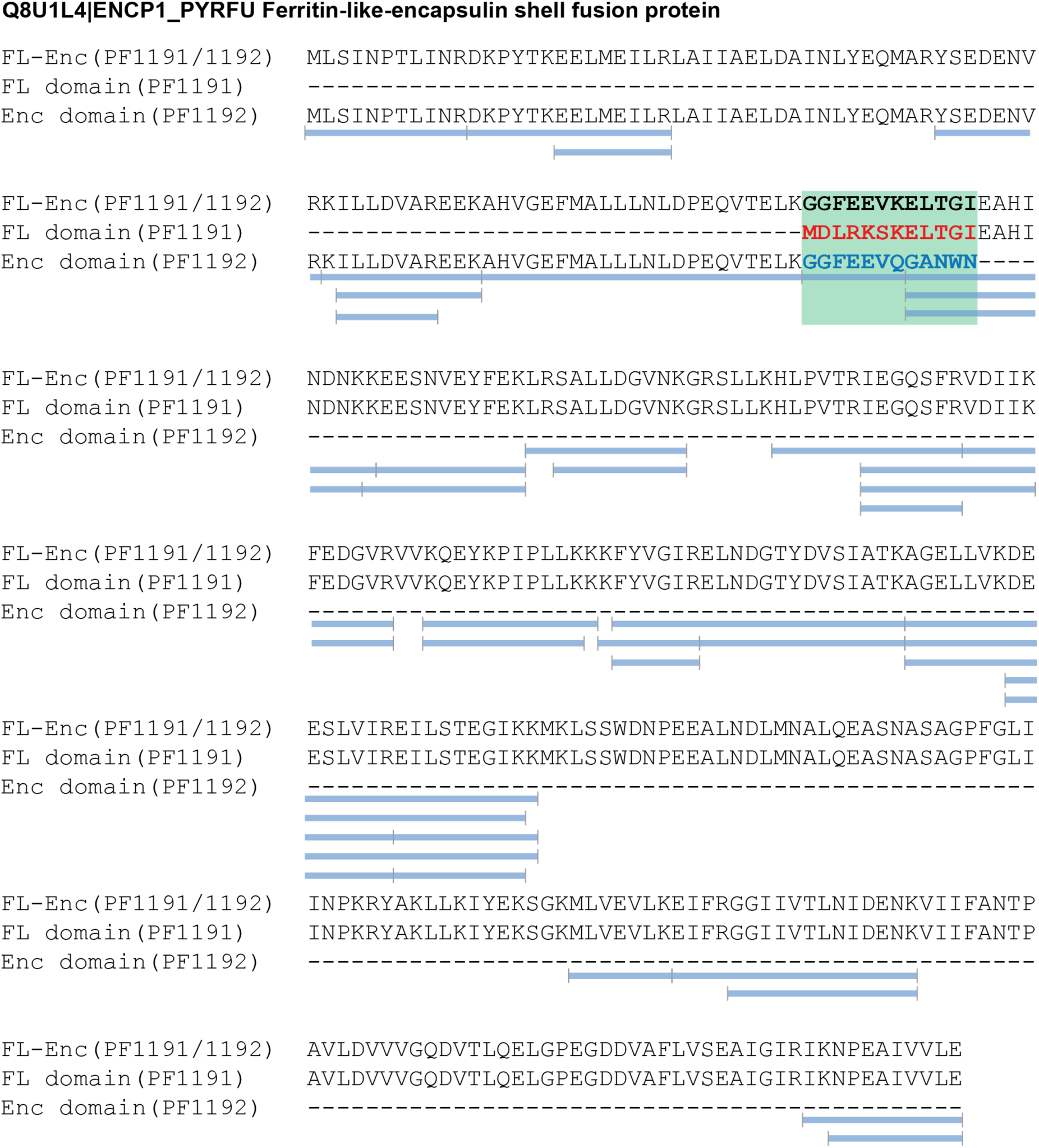
Sequence alignment and peptide mapping of Ferritin-like encapsulin shell fusion protein. Alignment of separate Enc (PF1192), the FL domain (PF1191) and FL-Enc fusion p<u>rotein</u> shows specific sequence interconnecting FL and Enc domains. Tryptic peptides identified by bottom-up MS (blue bars) yielded 64.9% sequence coverage, covering unique peptide sequences (green bar) responsible for fusion of FL and Enc domains.

**Supplementary Fig. 9.**
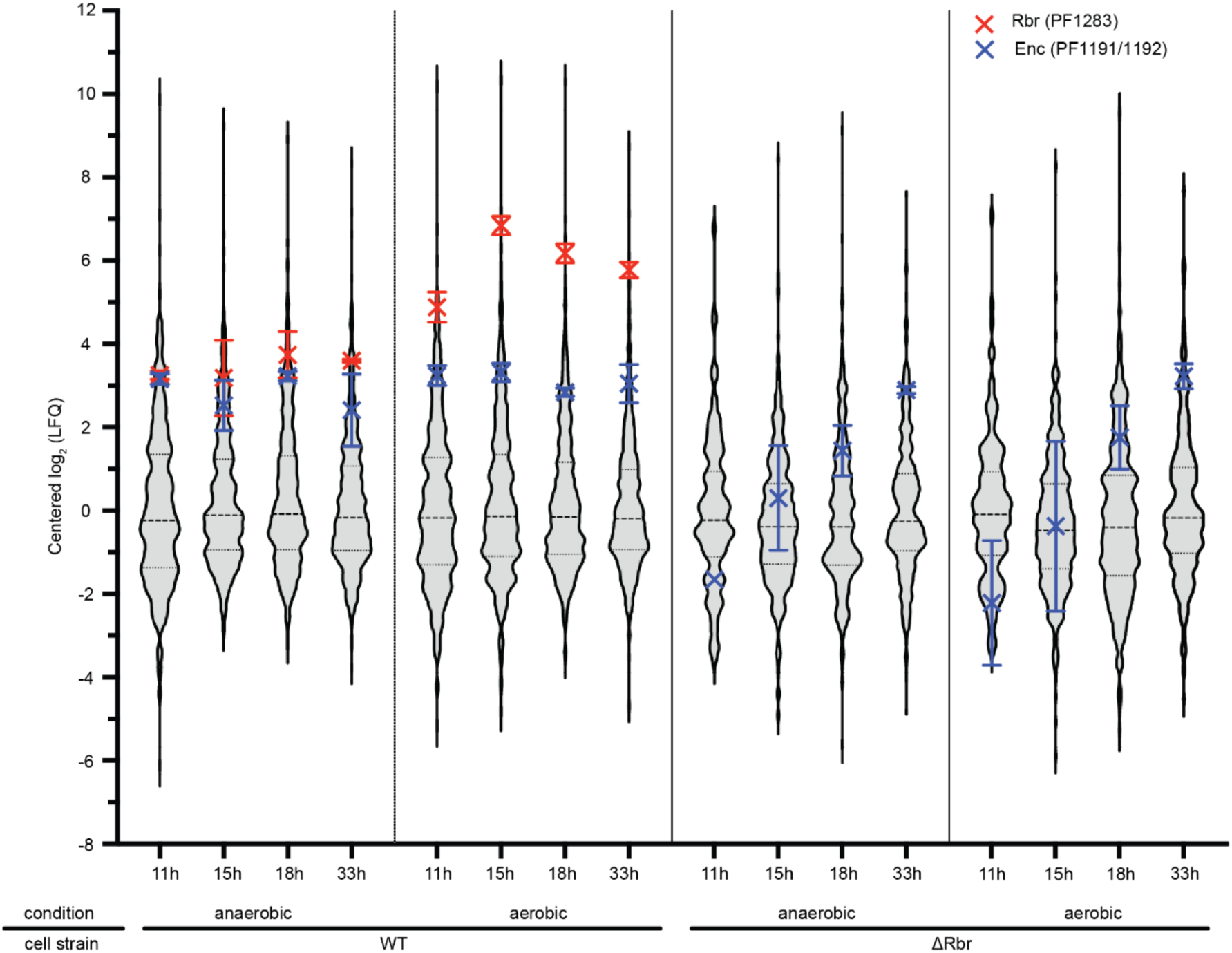
Proteome changes in *Δrbr* cells. Violins indicate changes of all proteins at different timepoints. Rbr can only be detected in the WT cells, consistent with the knock-out. Enc, which serves as a control, is detected in both cell types and does not change significantly.

**Supplementary Fig.10.**
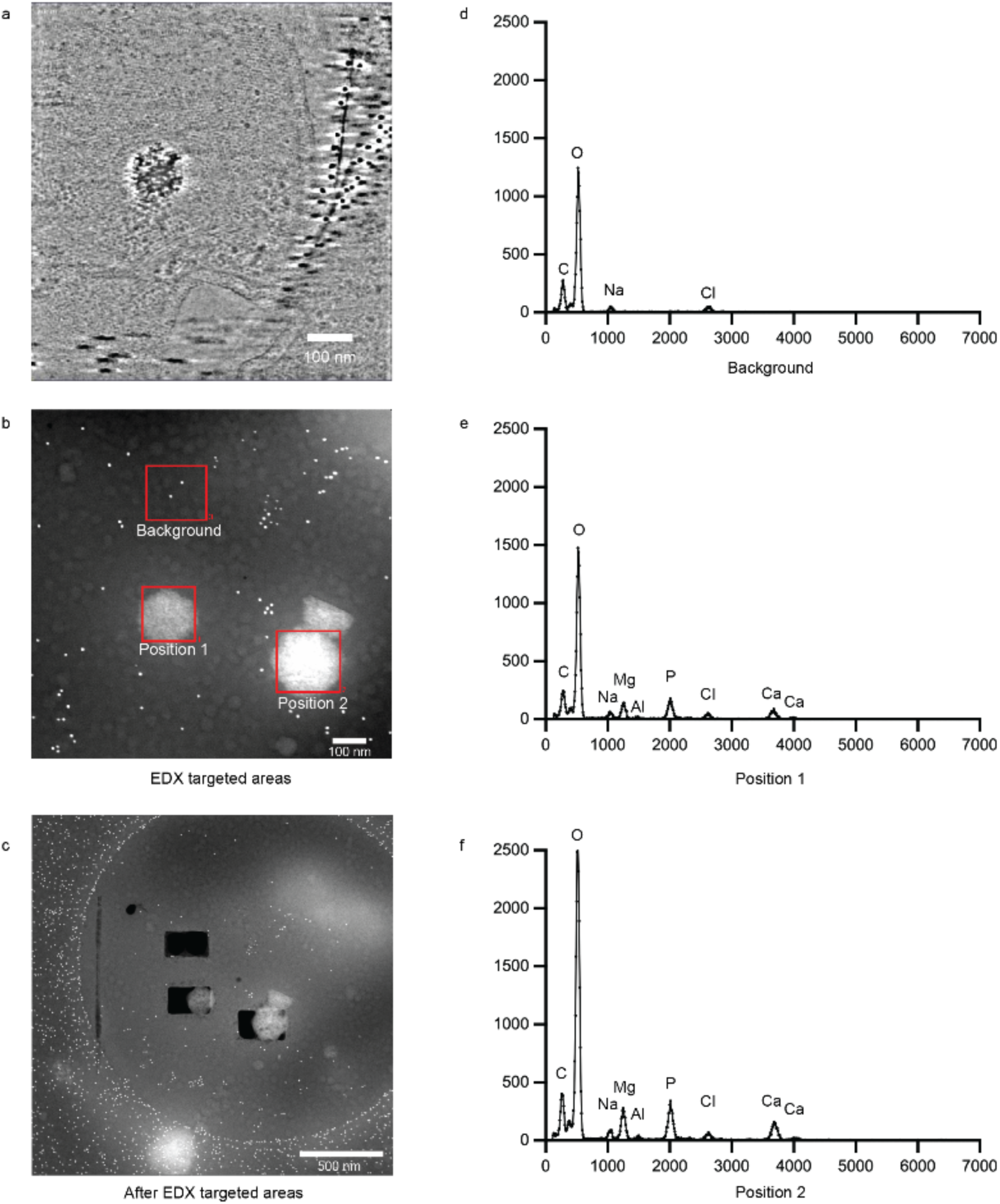
Influence of rbr knockout on *P. furiosus* cells. (a) Cryo-EM image of the oxidative stressed *Δrbr* cell shows intense inorganic aggregates inside the cell. Scale bar: 100nm. (b) High Angle Annular Dark Field (HAADF) cryo-EM image of the oxidatively stressed *Δrbr* cells used for EDX scanning. Two positions with strong signal likely arising from inorganic aggregates were selected for EDX, as well as a control area without strong HAADF signal. Scale bar: 100nm. (c) HAADF cryo-EM images of (b) after EDX scanning, indicating removal of vitrified water and cell organic components by the electron beam. Scale bar: 500nm. (d-f) EDX spectra in the areas marked “background”, “1” and “2” in b.

## Tables

**Supplementary Table 1.** CryoEM data collection and image processing statistics.

**Supplementary Table 2.** Data for volcano plots at 11h, 15h, 18h and 33h of culture growth time.

## Supplemental Movies

**Supplementary Video 1.** Cryo-electron tomogram of *P. furiosus* cell from culture grown for 33 h under anaerobic conditions.

**Supplementary Video 2.** Cryo-electron tomogram of *P. furiosus* cell from culture grown for 33 h under anaerobic conditions then exposed to oxygen containing environment.

**Supplementary Video 3.** 3D isosurface rendering of segmented cryo-electron tomogram of *P. furiosus* cell shown in supplemental movie 2.

**Supplementary Video 4.** Another example of cryo-electron tomogram of *P. furiosus* cell from culture grown for 33 h under anaerobic conditions then exposed to oxygen containing environment.

**Supplementary Video 5.** 3D isosurface rendering of segmented cryo-electron tomogram of *P. furiosus* cell shown in supplemental movie 4. Different from Supplementary movie 3, ribosomes are also shown in green.

**Supplementary Video 6.** Cryo-electron tomogram of Δrbr *P. furiosus* cell from culture grown for 33 h under anaerobic conditions then exposed to oxygen-containing environment, revealing dense aggregates.

## Methods

*P. furiosus* Cell cultivation and cultivation growth curve *P. furiosus* (strain: DSM3638, RRID: NCBITaxon_186497) cell inocula were obtained from the in-house culture collection at Wageningen University. Cells were grown with 250 ml medium in a 500-ml serum bottle at 95 °C for 33 h, with 0.5 atm overpressure with N_2_. Cells were collected every h after 7 h growth to monitor optical density at 600nm.

1 liter *P. furiosus* medium composition: KH_2_PO_4_ 0.14 g, CaCl_2_ 0.14 g, NH_4_Cl 0.25 g, MgCl_2_•6H_2_O 2.7 g, MgSO_4_•7H_2_O 34 g, KCl 3.3 g, 1 g NaHCO_3_, 25 g NaCl, 0.5 g L-Cysteine•HCl, 1 ml of 1000x NaWO_4_ (10 mM) solution, 1 ml of 1000x trace elements solution (2 g/l FeCl_2_.4H_2_O, 0.05 g/l H_3_BO_3_, 0.05 g/l ZnCl_2_, 0.03 g/l CuCl_2_, 0.05 g/l MnCl_2_•4H2O, 0.05 g/l (NH_4_)_6_Mo_7_O_24_.4H_2_O, 0.05 g/l AlCl_3_, 0.05 g/l CoCl_2_•6H_2_O, 0.05 g/l NiCl_2_, 0.5 g/l EDTA, 1ml HCl), 1 ml resazurin solution (5 g/l), 1 ml of 1000x vitamin mix solution (2 mg/l Biotin, 2 mg/l Folic acid, 10 mg/l Vitamin B_6_ (Pyridoxin HCl), 5 mg/l Vitamin B_2_ (Riboflavine), 5 mg/l Vitamin B_1_ (Thiamin HCl), 5 mg/l Nicotinamide, 5 mg/l Vitamin B_12_ (Cyanocobalamin), 5 mg/l p-Aminobenzoic acid, 5 mg/l Lipoic acid (Thioctic acid), 5 mg/l Pantothenic acid) and 40 mL of 25x amino acid solution (5.0 g/l each of glutamic acid and glycine, 3.125 g/l each of arginine and proline, 2.5 g/l each of asparagine, histidine, isoleucine, leucine, lysine, and threonine, 1.875 g/l each of alanine, methionine, phenylalanine, serine, and tryptophan, 1.25 g/l each of aspartic acid, glutamine, and valine and 0.3 g/l tyrosine). When necessary, the medium was supplemented with 20 μM uracil to allow for the growth of the ΔPyrF COM1 strain. As carbon source, we used 5 g/l maltose.

### Oxygen-stress treatment of *P.furiosus*

Cells were initially grown under anaerobic conditions. For the anaerobic conditions, cells were handled in an anaerobic hood and all experiments were conducted within the hood. To induce oxygen stress, after a specific cultivation period, the cells were exposed to air for at least 30 minutes until the resaurin in the culture changed from transparent to pink, confirming oxidation. The cells were then processed under aerobic conditions.

### Samples preparation for whole-cell mass spectrometry

Cells grown under anaerobic and oxygen-stressed conditions were collected at the time points of 11, 15, 18, and 33 h, pelleted, and the supernatant was removed. All time points were collected in triplicate. Subsequently, 30 μL of lysis buffer (100mM TRIS pH 8.5, 10mM TCEP, 40mM CAA, 1% sodium deoxycholate (SDC), supplemented with cOmplete Mini protease inhibitor (Roche) was added to each sample followed by heating to 95°C for 5 min. Samples were spun down and lysed by a Bioruptor plus (Diagenode) sonicator for 15 cycles (30 s on, 30 s off) set to intensity level 5. After lysis cell debris was spun down for 15 min at 20,000g and the supernatant was pipetted to new microtubes. Next, Benzonase® Nuclease (Merk) was added to a final concentration of 1 % and samples were incubated for 2 h at room temperature prior to vortexing at 800 RPM. After incubation, the concentration of extracted proteins has been determined by BCA Protein Assay Kits (Thermo Fisher Scientific). For digestion, a volume corresponding to 1 μg of protein was pipetted into new microtubes, and 50 μl of digestion buffer (200mM 4-ethylmorpholine, pH 8.5) was added. Subsequently, samples were digested overnight at 37 °C by trypsin (1:25 enzyme to protein ratio, Promega) and LysC (1:50 enzyme to protein ratio, Wako). Lysis was stopped by the addition of TFA (final concentration 2 %) which caused acetic precipitation of SDC. Samples were spun down at 20,000g for 20 min and peptides in supernatant were desalted on Oasis HLB 96-well µElution Plate (Waters Corp) using the vendor-provided protocol, followed by air drying on a vacuum centrifuge (Eppendorf). Dried samples were stored at -20°C prior to MS analysis.

### LC-MS/MS data acquisition and processing

Before analysis, dried peptides were reconstituted in solvent A (0.1% FA in water) and analyzed on UltiMate 3000 RSLCnano System (Thermo Fisher Scientific) coupled to Orbitrap Exploris 480 mass spectrometer (Thermo Fisher Scientific). For analysis, the equivalent of 0.5 µg of peptides was injected and desalted on trapping column Acclaim PepMap 100 C18 (5 mm × 0.3 mm, 5 μm, Thermo Fisher Scientific) in 0.1% FA at a flow rate of 30 μL/min followed by separation on in-house prepared analytical column (50 cm × 75 µm, 2.4 µm, ReproSil-Pur 120 C18Aq, Dr. Maisch HPLC) with flow rate set to 300 nL/min. Trap and analytical columns were kept at 40 °C in built-in oven. Peptides were separated in a gradient of solvent B (0.1% FA in 80% ACN) starting at 9 % B followed by gradient steps: 1-2 min 9-13% B, 2-39 min 13-44 % B, 39-44 min 44-55 % B, 44-45 min 55-99 % B, 45-50 min 99-99 % B, and 50-60 min 99-9 % B. MS data were recorded in data-dependent acquisition mode, with MS1 scans (one MS scan per second) recorded in a mass range of 375-1600 m/z at a resolution of 60,000, standard AGC target, and automatic injection time control. For MS/MS, the most intense precursor ions were selected and isolated by a quadrupole filter with an isolation window set to 1.4 m/z. Subsequently, precursor ions were fragmented by HCD fragmentation with a normalized collision energy (NCE) of 28 %. Fragmentation spectra were recorded with an Orbirap resolution of 15,000, standard AGC target, and automatic injection time mode. Dynamic precursor exclusion time of 10 s was set between two MS1 scans.

Raw LC-MS/MS data were processed by MaxQuant version 2.4.2.0 utilizing the Andromeda search algorithm^20^. The search was performed against a FASTA file containing 6125 entries (506 Swiss-Prot and 5619 TrEMBL) accessed 23.05.2023 through UniProt Knowledgebase^21^ limited to taxon ID 2261 corresponding to *P. furiosus* COM1 and *P. furiosus* (strain ATCC 43587 / DSM 3638 / JCM 8422 / Vc1) strains. An integrated MaxQuant contaminant database was used to assign common contaminant proteins. For search, enzyme specificity was set to Trypsin/P (C-terminal cleavage of lysine and arginine) with up to 3 missed cleavages. The minimal peptide length was set to 5 and the maximal peptide mass was set to 6,000 Da. For peptide search, fixed carbamidomethylation of cysteine and up to 5 variable modifications per peptide were allowed, namely: methionine oxidation; lysine, and protein N-termini acetylation; asparagine and glutamine deamidation; serine, threonine, and tyrosine phosphorylation. For the database search, precursor and fragment ions mass error was set to 4.5 and 20 ppm, respectively. Match between runs was allowed within a retention time window of 0.4 min. For peptide quantification, the Label-Free Quantification (LFQ)^22^ feature was enabled requiring at least 2 peptides per protein to be presented for sample pair-wise comparison. A false discovery rate (FDR) of 1 % for peptide spectrum matches (PSMs) and proteins was used. The output result file from MaxQuant “proteinGroups.txt” was further processed by Perseus version 2.0.7.0^23^, and the final data were visualized by GraphPad Prism 9.

### Knockout of Rbr

To knock out the *Rbr* gene in *P. furiosus* we used an adapted strategy (Michael Terns & Ryan Catchpole, unpublished data) previously used in various *Sulfolobus* species^24^. This strategy involved the use of the endogenous *P. furiosus* CRISPR-Cas system^25^ to target and eliminate chromosomes which retained a wild type (WT) genotype. Briefly, a knockout cassette was constructed which contained: 1) 500 bp homology arms upstream and downstream of the *Rbr* gene to facilitate integration of the knockout cassette at the target locus; 2) a 250 bp pop-out region which has homology to the 3’ end of the upstream homology arm and is used to facilitate a crossover event and removal of the *pyrF* selection marker; 3) a *pyrF* selection marker to facilitate efficient selection on medium without uracil (selects for integration of the knockout cassette) and counterselection on medium with 5-FOA (selects for pop-out) and; 4) a mini CRISPR array (repeat-spacer-repeat) which encoded a spacer (5’-AGTATACAACAAAGCCGCAGAATTCCAAGGAGAAAAG-3’) targeting the *Rbr* gene.

The knockout cassette was transformed into *P. furiosus* COM1 (Δ*pyrF*) following previously established protocols ^26^. After transformation and selection on media without uracil (selecting for the integration of the *pyrF* gene at the target locus), individual colonies were screened through PCR (Fw primer: 5’-AGATGAGGAAGCATTATTTTACC-3’ and Rv primer: 5’-AAGAGCATAGAGCTTGCC-3’) for the detection of the three potential genotypes (WT, *Δrbr* and *Δrbr*+*pyrF*). Colonies that showed either the *Δrbr* or the *Δrbr+pyrF* genotype, were grown on solid medium containing uracil and 5-FOA to facilitate the selection of pop-out mutants (i.e., *pyrF* is removed from the genome). Resulting colonies were screened again through PCR to confirm the presence of *Δrbr* mutants. *Δrbr P. furiosus* cultures were used to create anaerobic glycerol stocks and stored at -80oC until further use.

### OSIT enrichment

*P. furiosus* cell mass was obtained from Archaea Center Regensburg (Institute of Microbiology and Archaea Centre at the University of Regensburg). Cells (0.5-0.8 g) were defrosted and resuspended in 1 ml lysis buffer (10mM Hepes-NaOH pH 7.4, 150 mM NaCl, 10% wv sucrose, 5 mM MgCl2, protease inhibitor cocktail (Roche)) and lysed with a 2ml dounce homogenizer (Bellco Glass, Inc) (Pestle A for 30 times, pestle B for 30 times, on ice). The lysate was pelleted at 600 g for 10 min, then the supernatant was gently loaded onto a 30%-60% sucrose gradient and ultracentrifuged at 32,000 rpm for 14 h (Optima XPN-80 Ultracentrifuge, BECKMAN COULTER). Samples were collected by a gradient collector (FC 203B, GILSON), and each fraction was checked by negative staining and EM for OSITs. Samples with high OSIT enrichment were merged, 10x volume of dialysis buffer was added (10 mM Hepes-NaOH pH 7.4, 150 mM NaCl, 5 mM MgCl2, protease inhibitor cocktail (Roche)), and pelleted at 20,000g for 20 min. The final enriched OSIT sample was resuspended with dialysis buffer and analyzed by protein mass spectrometry and cryo-EM single particle analysis.

### Samples preparation and data treatment for proteomic analysis of enriched OSIT

Two independently purified aliquots of approximately 5-8 µg enriched OSIT were tryptic digested on S-Trap spin columns (ProtiFi) according to manufacturer instructions. Data were acquired by injecting 200 ng of peptides on the same set-up as for the whole-cell MS analysis described above. Raw data were processed using the MaxQuant software^20^ version 2.0.1.0 with standard settings applied as described above. In brief, an allowed precursor mass deviation of 4.5 ppm and an allowed fragment mass deviation of 20 ppm were set. Cysteine carbamidomethylation was set as static modification, and methionine oxidation, N-terminal acetylation as variable modifications (maximum five modifications per peptide allowed). For approximate absolute abundance of individual proteins, the iBAQ values in Extended Fig. 1a were derived by the normalization of the summed peptide intensities by the number of theoretically observable peptides for a given protein.

### Cryo-EM grid preparation

*P. furiosus* wild-type or Rbr knockout cells were collected in an anaerobic hood. The cells were pelleted and resuspended in their medium, and 3.5 μl sample were applied onto glow discharged EM grids (Quantifoil R2/2 Cu 200 mesh) under anaerobic conditions. Grids were blotted from the backside for 3-5 s and plunged immediately into liquid ethane using a manual plunger. The same protocol was applied to oxidative stressed *P. furiosus* cells, with all the preparation in the air after the sample collection. From collection to cryogenic fixation the sample was exposed approximately 30-60 min to oxygen.

For OSIT single particle analysis, 3.5 μl of enriched OSIT was loaded onto glow-discharged EM grids (Quantifoil R2/2 Cu 200 mesh) and immediately blotted for 4s from the backside using a manual plunger before plunging into liquid ethane.

### Lamella grid preparation

*P. furiosus* wild-type cells were collected and pelleted, then resuspended in their medium, and 3.5 µL of cell suspension was applied on glow discharged EM grids (Quantifoil R2/2 Cu 200 mesh) and plunged frozen using a manual plunger. EM grids were clipped into Autogrid and loaded into Aquilos 1 FIB-SEM system (Thermo Fischer Scientific), where they were sputtered with an initial platinum coat for 10s followed by a 10s gas injection system (GIS), thinned with a Gallium ion beam with current 500pA, 300pA, 100pA then polished at 30pA.

### Cryo-EM tomography and single particle data acquisition

Tilt series were collected on a Talos Arctica (Thermo Fisher Scientific) equipped with a K2 summit direct electron detector and energy filter (Gatan) operated at an acceleration voltage of 200 kV. Tilt images were recorded as series of 7-9 frames at defocus level ranging from -3 μm to -8μm and magnifications resulting in effective pixel sizes of 1.72 Å and 2.17 Å. Tilt series were acquired using SerialEM^27^ using a grouped dose-symmetric tilt scheme^28^ covering a range of ±51° and an angular increment of 3°. The cumulative dose of a series did not exceed 120 e^-^ Å^-2^. In total, 75 tilt series of anaerobic cells were acquired, and 578 tomograms of oxidative stressed cells were recorded, including 118 lamella tomograms.

The OSIT single particle cryo-EM dataset was collected on a Titan Krios (Thermo Fisher Scientific) equipped with a K3 summit direct electron detector and energy filter (Gatan). The TEM was operated at an acceleration voltage of 300 kV with an effective pixel size of 0.55 Å and cumulative dose 50 e^-^ Å^-2^. In total, 2,372 images were collected in super resolution mode, using SerialEM^29^ with acquisition areas identified manually.

### DTT treatment of OSITs imaged by negative stain EM

To enriched OSITs sample equal volume of DTT treatment buffer (10 mM Hepes-NaOH pH 7.6, 150 mM NaCl, 5 mM MgCl2, 10 mM DTT) was added resulting in a final concentration of 5 mM DTT and incubated on ice for 40 min. As a control, enriched OSITs were added to an equal amount of a control buffer lacking DTT (10 mM Hepes-NaOH pH 7.6, 150 mM NaCl, 5 mM MgCl_2_) and incubated on ice for 40 min. 3 μl sample were pipetted onto glow discharged 400 mesh carbon coated copper grids (400 Mesh Copper 100, Van Loenen Instruments) for 30 s. The grids were blotted from the edge side for 5s and washed with volume control buffer 3 times, prior to staining with 2% uranyl acetate for 30 s. Negative staining EM dataset were collected on a Talos L120C (Thermo Fisher Scientific) operated at an acceleration voltage of 120 kV with a Ceta CMOS camera (Thermo Fisher Scientific) at an effective pixel size of 17.1 Å (magnification 8,500X) and 4.06 Å (magnification 36,000X), respectively.

### Helical Reconstruction of OSIT Single Particle Analysis

Movie frames were aligned using MOTIONCORR2/1.6.4^30^, and contrast transfer function (CTF) parameters were estimated using CTFFIND-4.1^31^. Helical reconstructions^32^ were performed in Relion-3.1.4^33^. Particles were picked manually for the start -end coordinates and extracted in a box size of 1,800 pixels (98.4 nm) and down-scaled to 300 pixels. Reference-free 2D classification was performed to exclude false positive detections, followed helical 3D-autorefine with helical twist 21.7°, rise 80.1 Å. Subsequently, 3D classification without image alignment was performed to access different OSIT classes and to select suitable classes for C17 symmetry expansion. Only the minimally shifted particles, namely those with stable alignments, were retained and re-extracted in 3 tetramers-by-3 tetramers patches. 3D classification without image alignment was applied for the re-extracted 3 by 3 patches with 4 classes. One class particles were selected and randomly split into two subsets, then each subset was reconstructed by relion_reconstruct command, then two individual maps were applied for postprocessing, resulting in a structure with a resolution of 7.4 Å (GS-FSC = 0.143).

### Patch OSIT Single Particle Analysis

The collected dataset of 2,372 movies was imported to cryoSPARC v3.2^34^ for downstream processing. All movies were corrected for beam-induced motion with the patch-Fmovement job and their CTF parameters were calculated via the patch-CTF job. After pre-processing, the final 3×3 patch derived from the helical reconstruction, described above, was projected and used for template-based particle picking, resulting in an initial particle set of 2,444,405 single-particle images. The picked particles were extracted with a box size of 1024 pixels and Fourier cropped to 128 pixels (bin8) to speed up classification. 2D classification and *ab initio* 3D reclassification into 4 classes followed to further refine the particle set and discard junk particles. Then, the particles from the two best classes were selected, leading to a particle set of 1,141,467 particles. This set was re-extracted with a box size of 512 pixels and Fourier cropped to 256 pixels (bin2), then 2D reclassified into 250 classes. 2DF classes containing misaligned or junk particles were discarded and a refined set containing 425,731 particles was used for homogeneous refinement, applying a C2 symmetry. Finally, the particles were symmetry expanded (C2) and a local refinement was employed, using the map derived from the homogeneous refinement as an initial model, reaching a nominal resolution of 3.32 Å (GS-FSC = 0.143).

### Model building and refinement

Four copies of an AlphaFold2-derived model of the *P. furiosus* Rbr tetramer (UniProt ID: Q9UWP7) were placed into the EM-map and individually fitted in ChimeraX v1.7^35^. The resulting model and map were then used as input in Phenix v1.21^36^ for real-space refinement. Iron ions within the di-iron and RubL sites were transplanted from previously published crystal structures using the AlphaFill webserver^37^ and then manually refined in Coot v0.98^38^ . Iron coordination site geometry and occupancy were validated with the CheckMyMetal (CMM) webserver v2.1^39^ .

### Subtomogram averaging of the OSIT

Movies of individual projection images were motion-corrected and combined into stacks of tilt series with determined CTF parameters in Warp (1.0.9)^40^. The combined stacks were aligned in Aretomo 1.25^41^ . Particles were picked manually for the start-end coordinates in IMOD 4.10.29 ^42^ and AddModPts applied for periodic particle coordinates along the z axis by PEET 1.18^43^ then the particle coordinates were converted into a Warp recognizable format (star). Subsequentially, the determined particle coordinates were used for subtomogram reconstruction in Warp 1.0.9^40^ at a pixel size 6.90 Å. The 3D-autorefine and 3D classification without image alignment were performed in Relion 3.1.4^44^ .

### Energy dispersive X-ray (EDX)

EDX data were collected on a Talos F200X (Thermo Fisher Scientific) operated at an acceleration voltage of 200 kV, equipped with X-FEG and a Super-X G2 EDX detector, with probe current 1 nA. High annular dark field scanning transmission electron microscopy (HAADF-STEM) images were acquired by TIA (Transmission Electron Microscopy Image Analysis), EDX regions were imaged and analyzed in cell regions with and without visible granules by TIA.

